# Tnfrsf13b Variants Enhance IgG-Mediated Complement Responses to Promote Bacterial Clearance

**DOI:** 10.64898/2026.11.23.768670

**Authors:** Lwar Naing, Ian P Connell, Nicole E Guibord, Sarah Showers, Isabella K Liu, Ainsley Hellens, Charles Cascalho-Platt, Carly Kerr, Mayara Garcia de Mattos Barbosa, Tracy Jo Pasieka, Vanessa P Raab, Nancy Nagy, Jeffrey L Platt, Marilia Cascalho

## Abstract

*TNFRSF13B* encodes the TACI receptor, necessary for differentiation of B cells into plasma cells. *TNFRSF13B* is highly polymorphic, yet common non-synonymous variants including those that impair the receptor structure and/or signaling are retained in the population. Here we show that mice carrying the Tnfrsf13b A144E variant, the murine homolog of the human TNFRSF13B A181E variant associated with common variable immunodeficiency, effectively clear host-adapted *Citrobacter rodentium* (a murine model of pathogenic *E. coli* infection) despite nine-fold decreased bacteria-specific IgG compared to WT mice. IgG from *Tnfrsf13b* mutant mice exhibited reduced sialic acid, terminal galactose, fucose, and showed enhanced complement effector functions, including greater C1q binding and membrane attack complex (C5b) deposition. Results show that some Tnfrsf13b variants enhance complement-dependent antibody effector function, thereby identifying TACI as a regulator of IgG effector properties and providing a potential explanation for the evolutionary persistence of *TNFRSF13B* mutations.

## INTRODUCTION

Tumor Necrosis Factor Receptor Superfamily 13B (*TNFRSF13B)* is among the most polymorphic genes in mammals, with over 415 missense and 173 synonymous mutations, including several dominant-negative alleles recorded in gnomAD browser (v4.1.0) ^1^. Missense mutations in *TNFRSF13B* are found in 10%-20% of patients with common variable immunodeficiency (CVID) and selective IgA deficiency ^2,3^. However, most individuals carrying *TNFRSF13B* missense mutations that have been linked to altered receptor function show no signs of clinical immunodeficiency ^4^. Furthermore, population genetic analyses using gnomAD browser indicate that the *TNFRSF13B* missense mutations are not selected against, based on the ratio of observed to expected missense variation ^1^. What drives *TNFRSF13B* missense mutation retention in human populations is not known. The essential functions of the encoded receptor in host defense appear inimical to a deleterious role of many if not all the retained *TNFRSF13B* missense mutations.

*TNFRSF13B* encodes TACI (transmembrane activator and calcium modulator and cyclophilin ligand interactor), a key member of the TNF receptor superfamily that is necessary for B cell activation and plasma cell differentiation ^5,6^. Upon binding of its ligands APRIL (a proliferation inducing ligand) and BAFF (B cell activating factor) to the cysteine-rich domains of TACI, adaptor proteins such as CAML and TRAFs are recruited ^7^, leading to activation of downstream transcription factors including NF-κB ^7^. NF-κB signaling is essential for immunoglobulin class-switch recombination ^8^, antibody production ^9^, and maintenance of B-cell homeostasis ^10^. Consistent with TACI’s essential functions in immunity, TACI-KO mice had decreased IgG, IgM and IgA^11^ and failed to produce antibodies against encapsulated bacteria^12^. In contrast, however, TACI-KO mice generated high-affinity antibodies against antigens from *Citrobacter rodentium (C. rodentium)*, accelerating bacterial clearance ^13^. These findings led us to hypothesize that some TNFRSF13B variants may confer selective advantages, such as enhanced antibody effector functions despite producing less antibodies.

Since most TNFRSF13B variants do not cause complete TACI-deficiency and are expressed in heterozygotic form, we investigated how common variants of Tnfrsf13b impact bacterial clearance using a murine model expressing the murine equivalent (A144E) of a common human TNFRSF13B variant (A181E) associated with CVID and IgA-deficiency. A study by Dong et. al reported that 10.42% of patients with CVID or selective IgA deficiency carried the A181E mutation compared to only 0.88% in the control healthy group ^3^. The A181E allele is observed in 0.5%-1% of the general population, with frequencies up to 2.5% in individuals with Finnish European ancestry^1^. In this study, we investigate the antibody responses in *Tnfrsf13b* A144E mutant mice following infection with host-adapted *C. rodentium*, a murine model for human enteric infections caused by attaching and effacing (A/E) pathogens, specifically enteropathogenic *E. coli* (EPEC) and enterohemorrhagic *E. coli* (EHEC).

We considered two alternate possible outcomes: 1. Expression of the A144E variant causes immunodeficiency and as such, increases susceptibility to host-adapted enterobacteria infection; 2. Expression of the A144E variant allows effective clearance in spite of decreased antibody production, owing to unique antibody properties that arise as a direct consequence of altered B cell development.

## RESULTS

### Tnfrsf13b mutant mice are susceptible to host-adapted *C. rodentium*

We previously showed that mice expressing a Tnfrsf13b A144E variant are resistant to infection by laboratory-grown, non-virulent *C. rodentium*^14^. Results showed that *C. rodentium* co-opts secretory IgA (sIgA) to activate the locus of enterocyte effacement to promote expression of *C. rodentium* virulence factors and establish productive infection. Because *Tnfrsf13b* mutant mice have impaired production of sIgA compared to WT mice, infection by non-virulent *C. rodentium* does not result in a productive infection. We confirmed that sIgA is the driving factor for resistance to non-virulent *C. rodentium* inoculum because polymeric immunoglobulin receptor-KO (pIgR-KO) mice that lack sIgA due to a deficiency in the transportation of IgA from blood to the mucosal surface, also displayed resistance to non-virulent *C. rodentium* (data not shown). Our findings are consistent with work from others showing that sIgA can be co-opted by *Escherichia coli, Lactobacillus, and Bifidobacterium* to increase adherence and colonization^6,15,16^.

Increased expression of virulence following host infection has been reported in numerous pathogens that include *Helicobacter pylori*^17^, *Shigella flexneri 2a*^18^*, Acrobacter butzleri*^19^, and *Escherichia coli (EHEC)* ^20^. In EHEC and *C. rodentium*, host adaptation results in hypervirulent conditions due to enhanced expression of Type Three Secretion System proteins and LEE-encoded regulator (Ler) ^21^ ^22^. In this study, we investigated if *Tnfrsf13b* mutant mice are susceptible to hypervirulent *C. rodentium* as it emerges from an infected host, in a model of host-to-host transmission. To this end, we infected C57BL/6 WT mice with 10^8^ CFU of non-virulent *C. rodentium* grown in the laboratory. At 7-day post inoculation, fecal samples containing host-adapted *C. rodentium* were collected and prepared as an inoculum. We then infected new mice cohorts, *Tnfrsf13b* mutants or WT mice, with 10^7^ CFU of the host-adapted *C. rodentium.* This host-adapted hypervirulent *C. rodentium* infection scheme is shown in Figure 1A.

**Figure 1:**
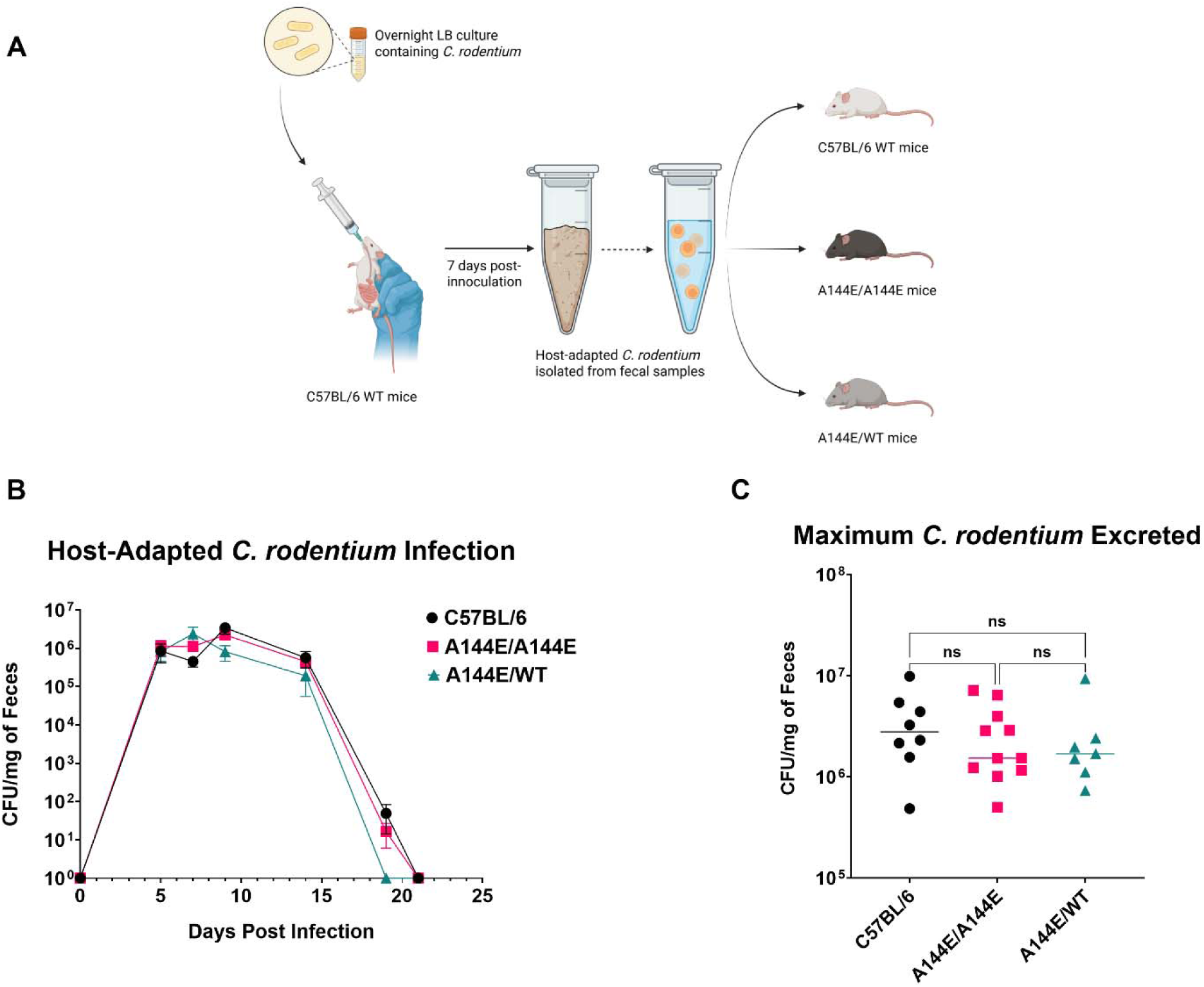
Host-adapted *C. rodentium* infection in *Tnfrsf13b* mutant mice. (A) Schematic depicts the infection of 8–12-week-old, age- and sex-matched WT and *Tnfrsf13b* mutant mice with 10 CFU of host-adapted *C. rodentium* administered by oral gavage. (B) Graph depicts the kinetics of *C. rodentium* infection quantified by the mean number of viable *C. rodentium* in feces (expressed in log_10_ CFU/mg of feces) at various times after infection. (C) Peak bacterial burden (maximum log₁₀ CFU/mg feces) observed during infection for each genotype. WT (n = 8), A144E/A144E (n = 11), A144E/WT (n = 7). Statistical analysis was performed using ordinary one-way ANOVA in GraphPad Prism v10.4.1.

We observed that both WT and *Tnfrsf13b* mutant mice exhibited similar infection kinetics following infection with host-adapted *C. rodentium*, with the peak of the infection occurring between days 7-10 post-infection (Figure 1B). Maximum fecal bacterial load did not differ between WT and *Tnfrsf13b* mutant mice (Figure 1C) and both WT and *Tnfrsf13b* mutant mice effectively cleared the infection by 21 days post inoculation (Figure 1B).

To determine if the susceptibility of Tnfrsf13b mutant mice to host-adapted *C. rodentium* reflects overcoming sIgA-mediated resistance or instead reflects properties of immunity that uniquely depend on the *Tnfrsf13b* genotype, we infected pIgR-KO mice shown previously to be resistant to infection with laboratory grown *C. rodentium*. Results show that pIgR-KO mice were as susceptible to host-adapted *C. rodentium* as WT mice, and the two groups had comparable infection kinetics and maximal fecal bacteria loads (Supplemental Figure 1). These findings indicate that sIgA-mediated resistance to laboratory-grown *C. rodentium* is overcome by host-adapted bacteria and that *Tnfrsf13b* mutant and pIgR-KO mice clear the host-adapted infection as well as WT mice.

### Tnfrsf13b mutant mice effectively clear *C. rodentium* despite decreased bacteria-specific antibodies

Clearance of *C. rodentium* and host survival requires adaptive immunity, and IgG in particular ^23,24^. While exact mechanisms to how bacteria-specific IgG is induced and reach gut lumen are unclear, systemic IgG localizes to the gut lumen to promote bacteria clearance. IgG is thought to migrate into the lumen by neonatal Fc receptor (FcRn)-dependent epithelial transport or by passive transfer following infection-associated disruption of epithelial junctions causing increased gut permeability ^25,26^. Epithelial barrier disruption also allows systemic exposure to *C. rodentium* antigens to facilitate the development of adaptive immune responses ^27,28^.

Ghaem-Maghami et. al^29^ showed that subcutaneous immunization with intimin, a key *C. rodentium* virulence factor, generated intimin-specific IgG responses that conferred protection against oral challenge with live *C. rodentium*, establishing intimin as an effective target for acquired immunity. To evaluate the adaptive immune responses to *C. rodentium* in *Tnfrsf13b* mutant mice, we measured serum *C. rodentium*-specific IgG including intimin-specific IgG and *C. rodentium* lipopolysaccharide (LPS)-specific IgG throughout the infection.

We found that in comparison to WT mice, *Tnfrsf13b* mutant mice produced significantly less *C. rodentium* specific IgG, intimin-specific IgG, and LPS-specific IgG following infection. (Figure 2A-C). At 19 days post-infection, mono and biallelic Tnfrsf13b mutant mice exhibited, on average, 5-fold and 8.3-fold less intimin-specific IgG, respectively, compared to WT mice (mean of 50 µg/mL in WT vs 10 µg/mL in A144E/A144E vs 6 µg/mL in A144E/WT) (Figure 2A).

**Figure 2:**
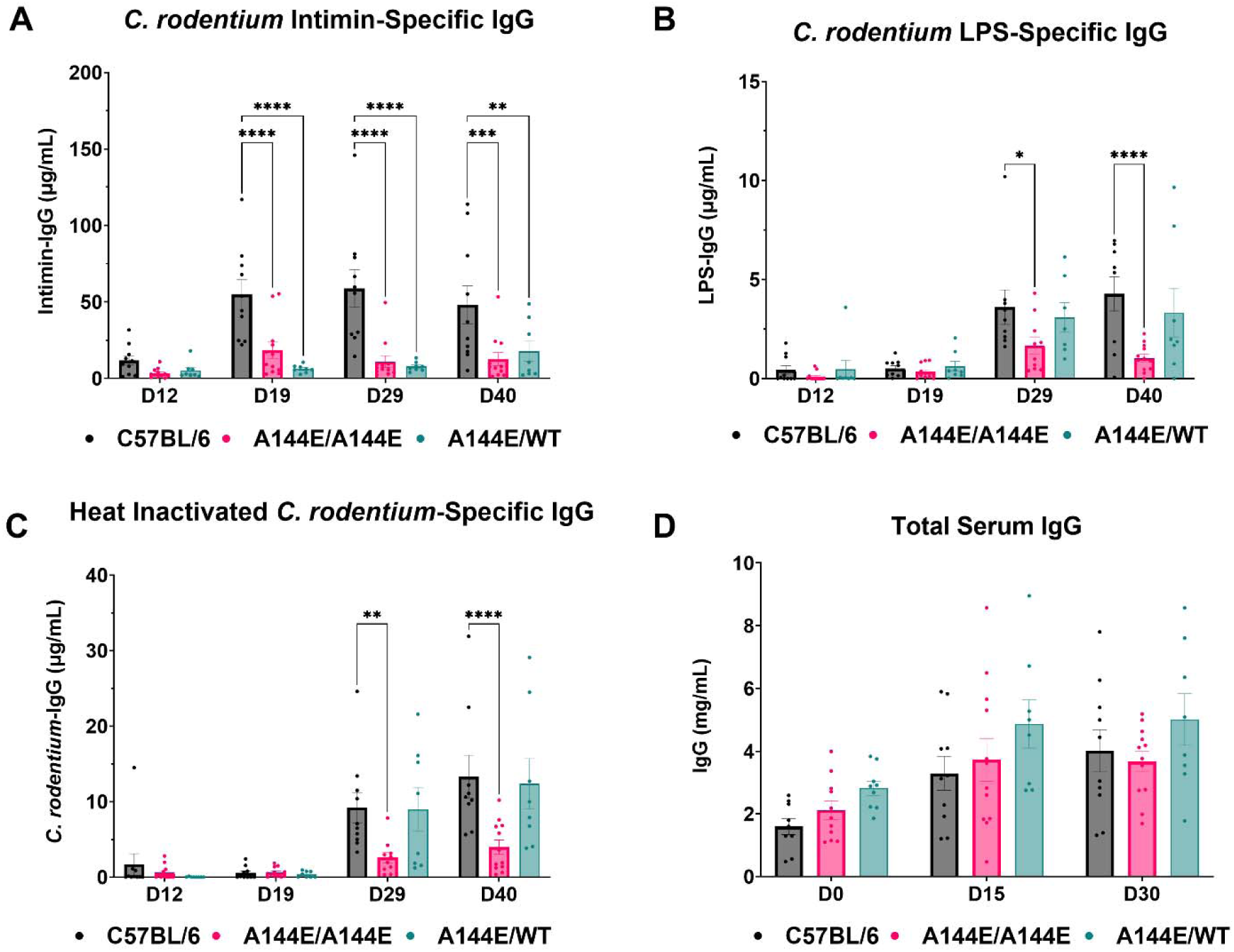
*Tnfrsf13b* mutant mice produce less *C. rodentium* specific IgG. Serum from infected mice were collected at indicated times following infection. Concentrations of (A) intimin-specific IgG, (B) LPS-specific IgG, (C) heat-inactivated *C. rodentium* specific IgG, and (D) total serum IgG were measured by ELISA. WT (n = 10), A144E/A144E (n = 12), A144E/WT (n = 8). Bar graphs show mean IgG concentrations. Statistical comparisons between genotypes are made with two-way ANOVA followed by Fisher’s LSD (Least Significant Difference) test in GraphPad Prism v10.4.1.

By 29 days post-infection, the difference in *C. rodentium-*specific IgG between WT and *Tnfrsf13b* mutant mice increased. *Tnfrsf13b* mutant mice had, on average, 8.6-fold less intimin-specific IgG, compared to WT mice (mean of 60 µg/mL in WT vs approximately 7 µg/mL in *Tnfrsf13b* mutant mice) (Figure 2A). Because *Tnfrsf13b* mutant mice displayed a decline in the concentrations of intimin-specific IgG between days 19 and 29 post infection, we also determined the overall protective immunity by rechallenging the mice with a larger inoculum, containing 10^8^ CFU of host-adapted *C. rodentium* 40 days after the initial infection. None of the mice experienced reinfection (data not shown), indicating that immunity generated in *Tnfrsf13b* mutant mice is protective and confers memory despite reduced concentrations of *C. rodentium*-specific IgG compared to WT mice.

We measured total serum IgG before and after infection and saw no differences between WT and *Tnfrsf13b* mutant mice, indicating that impaired bacteria-specific IgG responses in *Tnfrsf13b* mutant mice are not due to hypogammaglobulinemia (Figure 2D). Moreover, we did not detect *C. rodentium*-specific IgG antibodies in the fecal supernatant after infection in either WT or *Tnfrsf13b* mutant mice (data not shown), indicating that protection might be manifested at the mucosal and submucosal surfaces ^23^.

To check whether IgM might contribute to early defense against *C. rodentium,* we quantified *C. rodentium*-specific serum IgM at day 12 post infection and saw that WT and *Tnfrsf13b* mutant mice produced comparable *C. rodentium*-specific serum IgM (Supplemental Figure 2). We further measured total sIgA in feces during infection and confirmed prior findings^14^ showing reduced fecal sIgA in *Tnfrsf13b* mutant mice 19 days post-infection compared to WT mice. The results indicate that *Tnfrsf13b* mutant mice effectively clear host-adapted *C. rodentium* and mount effective memory responses, despite producing significantly less bacteria-specific IgG antibodies compared to WT mice.

### Tnfrsf13b mutant mice produce serum IgG with less sialic acid and terminal galactose

Prior studies have established that *C. rodentium*-specific IgG antibodies are essential for bacterial clearance ^23,24^. To assess the importance of *C. rodentium*-specific IgG antibodies for bacterial clearance in this study, we infected homozygous quasi-monoclonal (QM) mice, which cannot generate *C. rodentium* specific antibodies due to contraction of the primary B cell repertoire specificity to recognize the hapten-4-hydroxy-3-nitrophenylacetate ^30^. QM mice were unable to clear host-adapted *C. rodentium* until around 50 days post-infection, compared to WT mice which clear the infection by 21 days post-infection (Supplemental Figure 4A-B). As expected, QM mice produced hardly any intimin-specific IgG and LPS-specific IgG by 35 days post-infection compared to WT mice (Supplemental Figure 4C-D).

Given the importance of *C. rodentium*-specific IgG to clear *C. rodentium*, we investigated other immunological properties that could compensate for decreased bacteria-specific IgG antibodies in *Tnfrsf13b* mutant mice. We first asked if functional IgG avidity for *C. rodentium* is increased. Serum *C. rodentium* specific IgG antibodies were quantified by ELISA, and equivalent amounts of *C. rodentium* specific IgG from WT and *Tnfrsf13b* mutant mice were incubated with a range of titrated concentrations of GFP expressing *C. rodentium*. Using flow cytometry to quantify the IgG concentration required for 50% bacteria binding, we determined that it takes slightly higher amounts of IgG from *Tnfrsf13b* mutant mice to bind 50% of bacteria (139 ng for A144E/A144E, 156 ng for A144E/WT), compared to 95 ng for WT mice (Figure 3A). These findings suggest that IgG avidity to *C. rodentium* was not enhanced and probably decreased in *Tnfrsf13b* mutant mice, ruling out increased antibody avidity as a contributing factor to effective bacterial clearance despite decreased bacteria-specific IgG responses.

**Figure 3:**
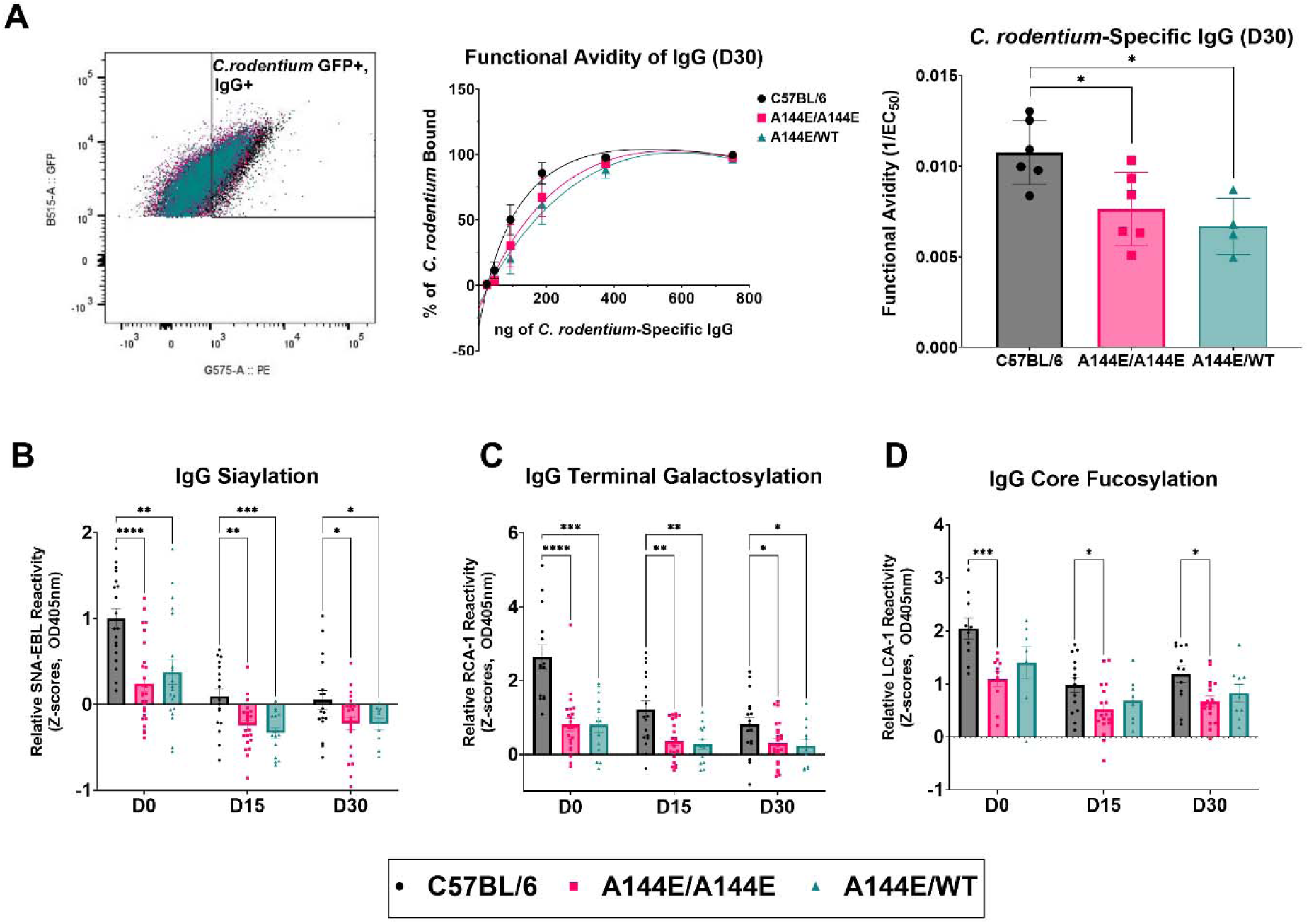
Serum IgG isolated from *Tnfrsf13b* mutant mice have less sialic acid, terminal galactose, and core fucose. (A) Functional avidity of IgG from WT (n = 6) and *Tnfrsf13b* mutant mice (A144E/A144E: n= 6, A144E/WT: n= 4) was assessed by incubating GFP expressing *C. rodentium* with equal amounts of *C. rodentium* specific IgG antibodies, quantified separately by ELISA. The fraction of IgG-bound GFP expressing *C. rodentium* was determined by flow cytometry; the gating strategy is shown in the left graph. The line graph shows the percentage of bacteria bound across increasing concentrations of IgG. The bar graph shows the amount of *C. rodentium*-specific IgG required for 50% bacteria binding, characterized as effective concentration (EC_50_). Functional avidity is reflected as (1/EC_50_). (B) IgG sialic acid, (C) terminal galactose, and (D) core fucose was measured in IgG isolated from serum from WT mice (B and C: n= 18, D: n= 11), bi-allelic A144E mice, (B and C: n= 23, D: n= 13), and mono-allelic A144E mice (B and C: n= 15, D: n= 8) at different times after infection. 25 µg of purified plasma IgG was used as the input. Lectin reactivity was quantified on IgG serial dilutions by OD405nm absorbance values by ELISA. Z-scores of lectin reactivity were calculated to mitigate the interplate variation. For IgG avidity, statistical comparisons between genotypes were made with ordinary one-way ANOVA followed by Tukey’s multiple comparisons test. Z-scores of lectin reactivity were analyzed by mixed-effects analysis followed by Fisher’s (LSD) tests. All statistical analysis was performed in GraphPad Prism v10.4.1.

We next examined the IgG glycosylation profile in *Tnfrsf13b* mutant mice. IgG Fc glycosylation is essential for regulating antibody effector functions, including Fc receptor engagement and complement activation. Loss of IgG N-glycans, including sialic acid and terminal galactose, is linked to increased activation of inflammatory responses in infections like SARS-CoV-2 ^31,32^ ^33^ or HIV-1 ^34^. Moreover, IgG antibodies without core fucose at Asn 297 glycan have increased affinity to Fc RIIIA expressed by natural killer (NK) cells and Fc RI expressed by macrophages ^35–37^. To evaluate the IgG glycosylation profile in WT and *Tnfrsf13b* mutant mice, we quantified specific glycan residues on purified IgG immobilized on ELISA plates, using biotinylated Ricinus communis agglutinin I (RCA-1), Sambucus nigra agglutinin (SNA-EBL), and Lens Culinaris Agglutinin (LCA) to detect terminal galactose (Galβ1-4GlcNAc), α2,6-linked sialic acid residues (Neu5Acα2-6Gal/GalNAc) on N-glycans, and α1-6 fucose attached to GlcNAc near asparagine respectively.

Prior to *C. rodentium* infection, purified serum IgG from both heterozygous and homozygous *Tnfrsf13b* mutant mice display significantly less sialic acid and terminal galactose compared to WT mice, as detected by lectin reactivity (Figure 3B-C). IgG core fucosylation was also decreased in homozygous *Tnfrsf13b* mutant mice (Figure 3D). Fifteen and 30 days after infection, sialic acid and galactose content of IgG decreased relative to measurements prior to infection in every mouse. The amount of sialic acid and terminal galactose on IgG remained lower in *Tnfrsf13b* mutant mice relative to IgG glycans isolated from WT mice. These results suggest that changes in the glycan composition of *Tnfrsf13b* mutant IgG may increase inflammatory responses which can aid in *C. rodentium* clearance.

### Plasma cells from Tnfrsf13b mutant mice produce less glycotransferases ST6GAL1 and B4GALT1

To investigate the mechanisms underlying reduced IgG sialylation and galactosylation in *Tnfrsf13b* mutant mice before and after the *C. rodentium* infection, we isolated splenocytes from naïve WT and from *Tnfrsf13b* mutant mice followed by culture for 72 hours in B cell media containing β-mercaptoethanol (BME) to support B cell stimulation and survival ^38^. Following 72 hours, we measured the expression of the sialyltransferase ST6GAL1 and the galactosyltransferase B4GALT1 in CD138⁺ plasma cells by flow cytometry. Both ST6GAL1 and B4GALT1 expression were reduced by more than 2.3 fold in plasma cells from *Tnfrsf13b* mutant mice in comparison to WT mice (Figure 4A and 4C).

**Figure 4:**
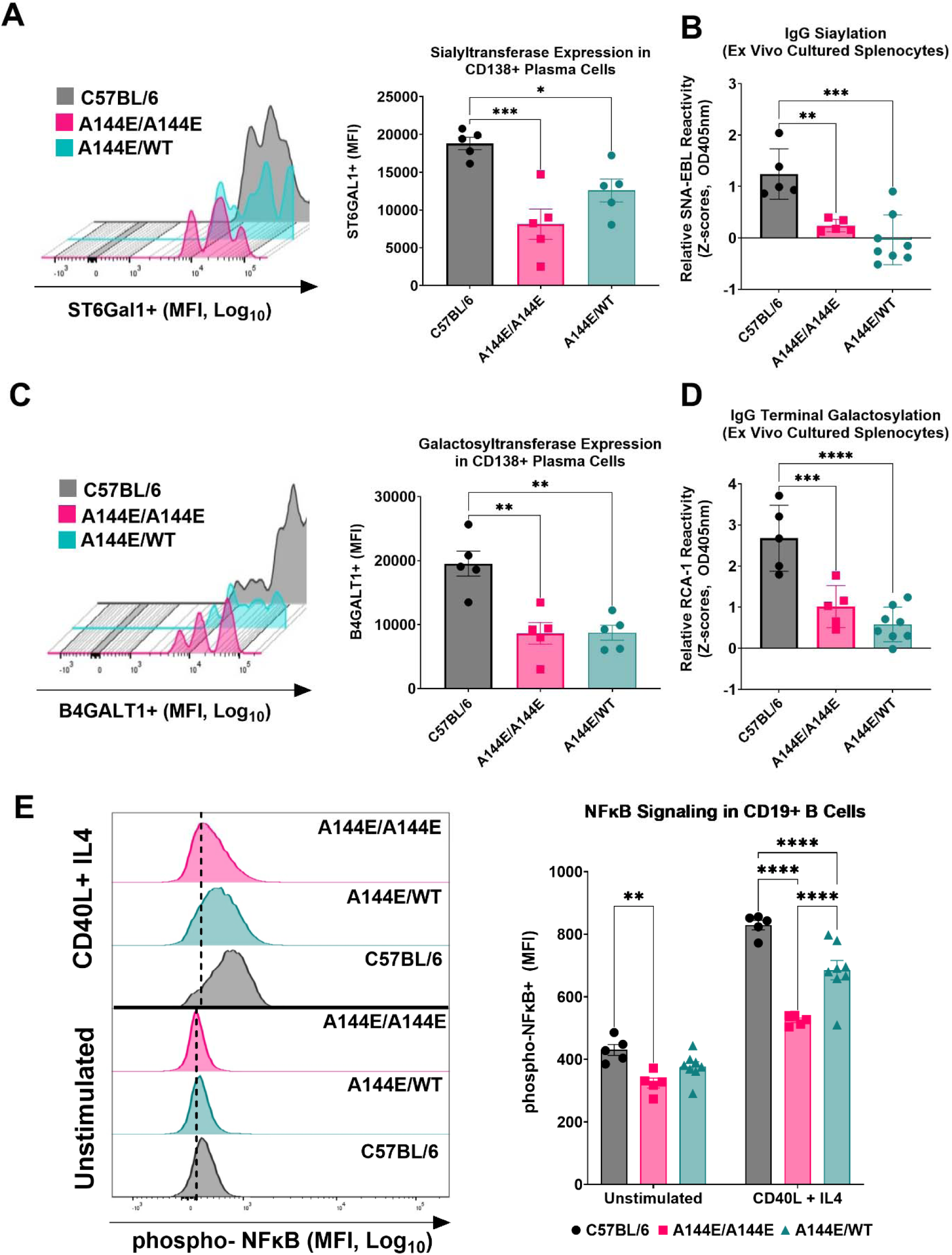
*Tnfrsf13b* genotype determines glycosyltransferase expression in plasma cells. (A) Expression of the sialyltransferase ST6GAL1 in CD138⁺ cells from WT and *Tnfrsf13b* mutant mice. Splenocytes were cultured for 72 hours in B cell media containing BME, then fixed, permeabilized, and intracellularly stained for ST6GAL1. CD138⁺ cells were gated for analysis. Representative MFI distribution curves and corresponding quantification (bar graphs) for WT, A144E/A144E, and A144E/WT mice (n = 5 per group) are shown. (B) IgG sialylation levels measured by lectin assay using IgG purified from cell culture supernatants of ex vivo–stimulated splenocytes. (C) Expression of the galactosyltransferase B4GALT1 in CD138⁺ cells following the same ex vivo culture and intracellular staining strategy described in (A). Representative MFI distribution curves and quantification are shown (n = 5 per group). (D) IgG terminal galactosylation assessed by lectin assay using purified IgG from culture supernatants. (E) Quantification of NF-κB signaling in splenic CD19+ B cells was quantified by isolating splenocytes from WT mice (n=5), A144E/A144E mice(n=5), and A144E/WT mice(n=8), and stimulating them with CD40L and IL4. Unstimulated and stimulated cells after 15 minutes of culture were fixed, permeabilized, and stained with anti-CD19 antibody and anti-phosphopeptide Ser536 NF-κB p65. Representative MFI distribution curves and corresponding quantification (bar graphs) for WT (n=5), A144E/A144E (n=5), and A144E/WT (n=8) are shown. Ordinary one-way ANOVA with Tukey’s multiple comparisons tests were used to analyze (A), (B), (C), and (D). Two-way ANOVA with uncorrected Fisher’s LSD test was used to analyze (E) on GraphPad Prism v10.4.1.

In parallel, we performed lectin-based ELISA assays to quantify IgG galactosylation and sialylation in the supernatant obtained from 72-hour B cell culture. Consistent with *in vivo* serum IgG glycosylation analysis, IgG from *Tnfrsf13b* mutant mice had less IgG galactose and less sialic acid compared to WT mice (Figure 4B and 4D).

To assess upstream signaling mechanisms that could link Tnfrsf13b signaling to altered glycosyltransferase expression, we evaluated canonical NF-κB activation by measuring phosphorylation of the p65 (RelA) subunit at serine 536 within splenocytes from WT and *Tnfrsf13b* mutant mice. Phosphorylation of p65 at Ser536 represents functional activation of NF-κB, corresponding to increased signaling, nuclear translocation, and transcriptional activity in B cells ^39,40^. Flow cytometry analysis showed that CD19+ B cells from *Tnfrsf13b* mutant mice produce less phosphorylated p65 constitutively and less phosphorylated p65 following 15 minutes of CD40L and IL-4 stimulation (Figure 4E). We used CD40L and IL-4 as stimulants for B cells because CD40L has been shown to activate both the canonical and non-canonical NF-κB pathways through interactions with TRAF adaptor proteins ^41,42^. Additionally, IL-4 has been reported to enhance NF-κB activation induced by other stimuli ^43–45^. TACI itself is a known activator of NF-κB, and CD40L, TACI, and TLR9 have been shown to synergize in driving B cell activation through convergent NF-κB signaling ^46^. In the absence of functional TACI signaling, this synergistic action is disrupted. Our results suggest that the A144E variant impairs synergistic potentiation of NF-κB activation in response to CD40L and IL-4 stimulation, resulting in reduced p65 phosphorylation in TACI variant B cells.

We also quantified Tnfrsf13b expression on splenic CD19+ B cells under unstimulated conditions and after 72 hours of CD40L and IL-4 stimulation. At baseline, *Tnfrsf13b* mutant mice exhibited increased Tnfrsf13b expression on splenic CD19+ B cells (Supplemental Figure 5). This is in contrast to findings from a prior study that analyzed B220+ splenic cells ^47^. However, following CD40L and IL4 stimulation, B cells from Tnfrsf13b A144E mice had reduced Tnfrsf13b expression on splenic CD19+ B cells compared to WT mice, suggesting that Tnfrsf13b expression is dynamically controlled in response to B-cell activation signals.

In addition to being important for canonical NF-κB signaling ^48^, CD40L and IL-4 ligands are commonly used to study T cell-dependent B cell responses^49^, which produce high-affinity antigen-specific antibodies against protein antigens such as intimin. The impairment in canonical NF-κB activation following CD40L and IL-4 stimulation in *Tnfrsf13b* mutant mice suggests that Tnfrsf13b, CD40, and other B cell receptors synergize to promote plasma cell differentiation and antibody-inducing transcriptional programs, consistent with previous findings^46,50^. Our findings thus support a model in which altered Tnfrsf13b signaling in *Tnfrsf13b* mutant mice reduces NF-κB activation and glycosyltransferase expression, ultimately changing IgG glycosylation profiles.

### *C. rodentium* isolated from infected Tnfrsf13b mutant mice have increased complement deposition

Based on our findings showing that *Tnfrsf13b* mutant mice have reduced sialic acid and galactose on IgG, we specifically tested whether those changes enhance complement activation following *C. rodentium* infection compared to WT mice. To assess complement activation *in vivo* during *C. rodentium* infection, WT and *Tnfrsf13b* mutant mice were infected with GFP expressing *C. rodentium.* Ten days after inoculation, we isolated GFP+ *C. rodentium* from fecal pellets extracted from the proximal region of the colon. Deposition of complement factors, C1q, C3d, and C3b, on the GFP-expressing *C. rodentium* were quantified by flow cytometry. GFP-expressing *C. rodentium* from *Tnfrsf13b* mutant mice exhibited increased C1q, C3d, and C3 deposition compared with bacteria from WT mice (Figure 5A-C). Notably, WT and *Tnfrsf13b* mutant mice had comparable bacterial loads (Figure 5D) and similar levels of IgG bound to the bacteria at 10 days post-infection (Figure 5E), indicating that enhanced complement deposition in A144E mice was not due to differences in bacterial load or IgG binding.

**Figure 5:**
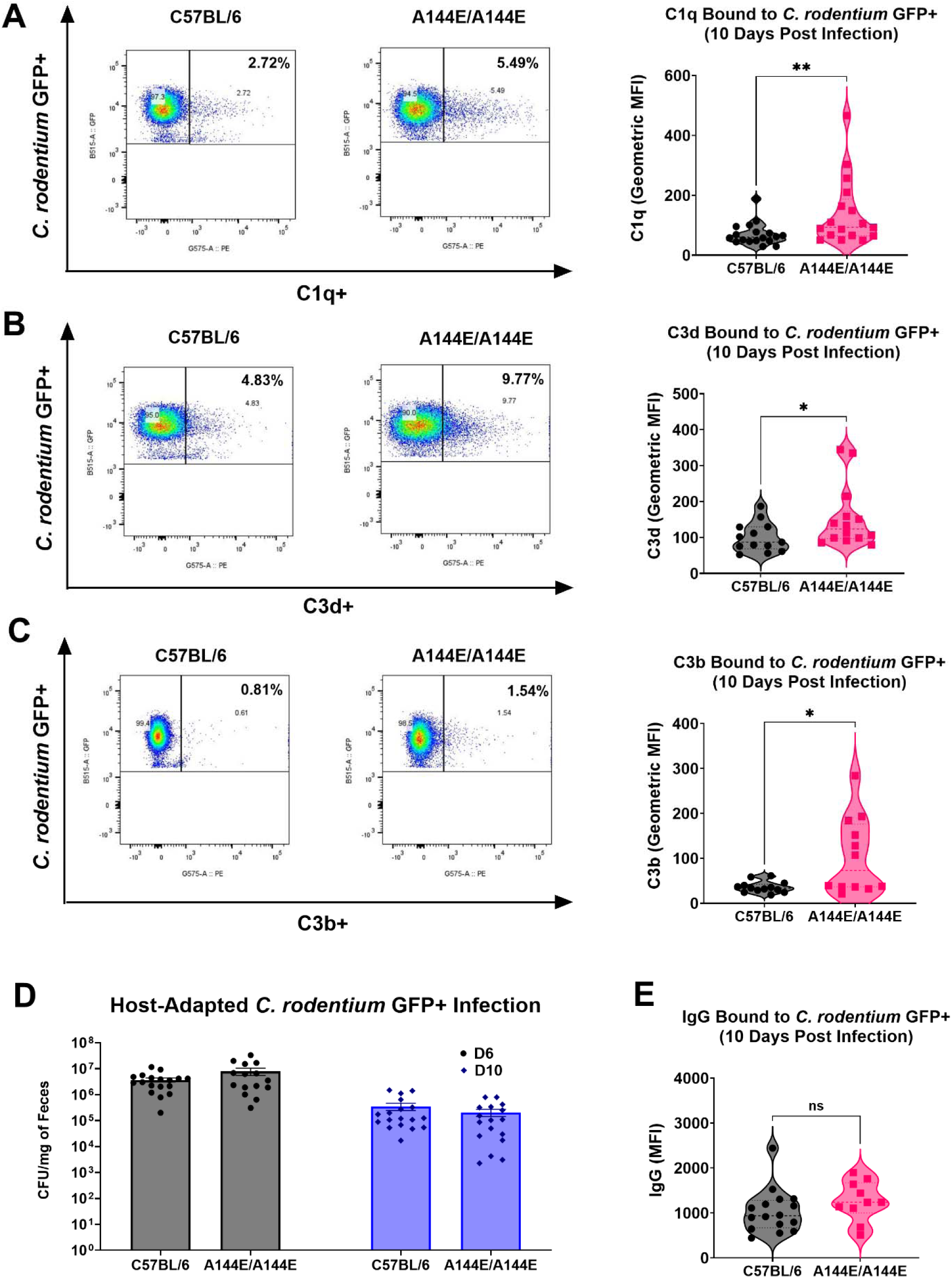
Complement deposition on *C. rodentium* isolated from infected WT and *Tnfrsf13b* mutant mice. Mice were infected with 10^7^ CFU of host-adapted, GFP-expressing *C. rodentium* via oral gavage. At 10 days post-infection, fecal pellets from the proximal colon were extracted and fecal bacteria were isolated. GFP-expressing *C. rodentium* was identified by flow cytometry. The amount of (A) C1q (WT: n= 18, A144E/A144E: n= 17), (B) C3d (WT: n=13, A144E/A144E: n= 12), (C) C3b (WT: n= 13, A144E/A144E: n= 12) deposition on GFP-expressing *C. rodentium* was measured. Representative flow cytometry plots showing complement deposition on GFP-expressing *C. rodentium* are shown, along with quantification of geometric mean fluorescence intensity (MFI) using violin graphs. (D) Mean and SE of viable *C. rodentium* in feces (expressed in log_10_ CFU/mg of feces) were measured at day 6 and day 10 post infection (WT: n=18, A144E/A144E: n=17). (E) IgG deposition on GFP-expressing *C. rodentium* quantified as MFI (WT: n= 16, A144E/A144E: n=10). Statistical comparisons between genotypes were made with Mann-Whitney Test in GraphPad Prism v10.4.1.

Sialic acid on IgG dampens inflammatory functions of IgG by limiting Fcγ receptor binding ^7,51^ and decreases TLR4 signaling and cytokine production ^52^. Quast et al. reported the direct inhibitory effect of IgG sialylation on C1q binding to galactosylated Fc fragments in polyclonal IgG ^53^. Our results indicate that reduced IgG core fucosylation, decreased IgG sialylation, and terminal galactosylation in Tnfrsf13b A144E variant mice may shift IgG towards more pro-inflammatory effector functions, including increased complement activation against *C. rodentium*.

### *C. rodentium* specific IgG isolated from Tnfrsf13b mutant mice exhibit enhanced complement recruitment in comparison to IgG isolated from WT mice

Given that *Tnfrsf13b* mutant mice exhibit increased complement deposition *in vivo,* we next asked whether purified bacteria-specific IgG antibodies from *Tnfrsf13b* mutant mice enhance complement activation. We quantified free serum C3 in WT and *Tnfrsf13b* mutant mice fifteen days after infection with *C. rodentium*. The free serum C3 levels were reduced in *Tnfrsf13b* mutant mice compared with WT mice (661.5 pg/mL vs 1233 pg/mL; Figure 6A), consistent with increased complement consumption during infection.

**Figure 6:**
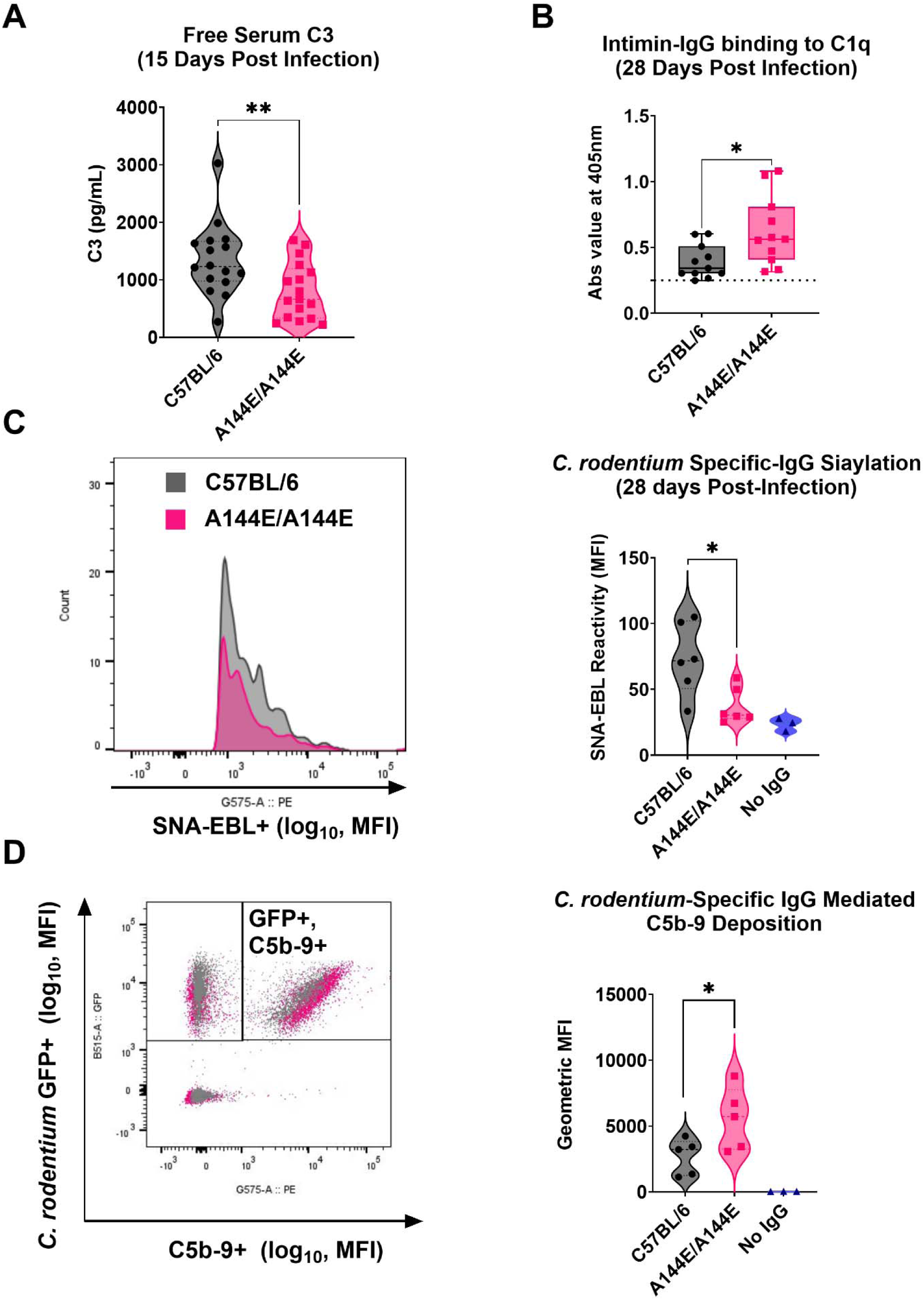
*Tnfrsf13b* mutant mice have increased complement consumption and enhanced complement fixation by *C. rodentium*-specific IgG. (A) Free serum C3 levels were measured at day 15 post infection for WT (n= 16) and Tnfrsf13b variant A144E mice (n=16) using Abcam C3 ELISA kit. (B) C1q recruitment was determined by incubating recombinant C1q protein with equivalent amounts of intimin-specific IgG from WT (n=11) and Tnfrsf13b variant A144E mice (n=11) at 28 days post infection. Bound C1q was quantified by anti-C1q ELISA using a plate reader. (C) *C. rodentium*-specific IgG sialylation was quantified by incubating GFP-expressing *C. rodentium* with normalized amounts of *C. rodentium*-specific IgG antibodies from WT (n=6) and Tnfrsf13b variant A144E mice (n=6) 28 days post-infection. SNA-EBL reactivity with bound IgG was analyzed by flow cytometry. Samples without IgG inputs were included to determine the background. Representative MFI distribution curve and quantification of SNA-EBL reactivity as mean fluorescence intensity (MFI) in a violin graph are shown. (D) C5b-9 deposition on GFP-expressing *C. rodentium* mediated by IgG isolated from WT (n=5) and Tnfrsf13b variant A144E mice (n=5) at 28 days post infection. The amount of IgG input in the assay is the equivalent amount of *C. rodentium*-specific IgG determined in a separate ELISA. Samples without any IgG were included to determine the background. Representative flow cytometry plots showing complement deposition on GFP-expressing *C. rodentium* are shown along with the quantification of geometric mean fluorescence intensity (MFI) using violin graphs. Statistical comparisons between genotypes were made with Mann-Whitney Test (A, B, and C) and unpaired T test (D) in GraphPad Prism v10.4.1.

To directly evaluate classical complement pathway initiation, we measured C1q recruitment by intimin-specific IgG isolated 28 days after infection. MaxiSorp plates were coated with intimin and incubated with equivalent amounts of purified IgG specific to intimin, followed by recombinant C1q. Bound C1q was quantified by ELISA using an anti-C1q antibody. We found that intimin-specific IgG from *Tnfrsf13b* mutant mice had higher C1q binding than equivalent amounts of intimin-specific IgG isolated from WT mice (Figure 6B), strengthening the concept that *Tnfrsf13b* genotype determines pathogen-specific IgG-mediated complement activation.

Because IgG sialylation has been shown to mediate C1q recruitment and downstream C3b deposition ^53^, we then specifically analyzed sialylation of *C. rodentium*-specific IgG. GFP-expressing *C. rodentium* was incubated with normalized amount of purified *C. rodentium* specific IgG from WT or *Tnfrsf13b* mutant mice 28 days after infection. We found *C. rodentium*-specific IgG from *Tnfrsf13b* mutant mice to have less sialic acid than WT mice (Figure 6C).

We next assessed complement activation downstream of IgG binding by measuring deposition of C5b-9. C5b-9, also known as the membrane attack complex (MAC), represents the terminal step of complement activation and mediates lysis of target cells or bacteria ^54^. Consistent with the increased C1q binding observed for *Tnfrsf13b* mutant IgG, *Tnfrsf13b* mutant IgG also mediated enhanced C5b-9 deposition on *C. rodentium* (Figure 6D). To confirm that complement activation in this assay was IgG-dependent and mediated through the classical pathway, we included control samples with an absence of IgG, containing only a complement source, which produced minimal signal.

Together, our data demonstrates that IgG-mediated complement activation against *C. rodentium* is enhanced in *Tnfrsf13b* mutant mice due to changes in the IgG glycosylation profile. Using a reductionist assay in which we analyzed purified IgG, bacteria, and complement in isolation, our results provide mechanistic evidence that *Tnfrsf13b* mutant mice can effectively clear *C. rodentium* infection despite producing less bacteria-specific IgG, owing to enhanced IgG-mediated complement activation.

## DISCUSSION

Antibodies produced by plasma cells are essential for bacterial clearance, including the clearance of enteric bacteria. Beyond preventing pathogen entry^55^, antibodies, specifically the Fc region of IgG, engage with the innate immune system, including natural killer cells^56,57^, macrophages^58,59^, and complement ^60,61^ to mediate pathogen killing. Activation of the classical complement pathway is initiated by the interaction between the Fc region of the pathogen-bound IgG and C1q to initiate the complement cascade that leads to pathogen killing^62,63^. Once C4 convertase is formed, the classical pathway helps activate the C3 convertase that is part of the alternative pathway^64^. Despite the importance of antibody-dependent complement activation in host defense, the genetic determinants that control antibody-mediated complement activation are not well defined.

*TNFRSF13B* is a highly polymorphic gene that governs many aspects of B cell differentiation and function, but how common variants contribute to host defense remains poorly understood. The results reported indicate that common polymorphisms known to impair TACI receptor function do not impair host defense despite reduced neutralizing antibody production. The finding is consistent with the concept that missense mutations in *TNFRSF13B* that impair receptor signaling^65^ and are associated with CVID and primary IgA deficiency enhance other immunological properties. Accordingly, we found that mutant *C. rodentium*-specific IgG activates complement more effectively, and in this way is able to enhance bacterial clearance^66^.

Our results link *TNFRSF13B* genotype to IgG glycosylation and control of inflammatory properties of IgG. Thus, the *Tnfrsf13b* genotype governs the amount of sialic acid, terminal galactose and fucose on IgG, in turn determining IgG engagement of complement. Terminal galactose is required for IgG Fc hexamerization following binding to antigenic targets, promoting C1q engagement ^67,68^. Galactosylated IgG also inhibits complement activation by facilitating interactions between FcyRIIB and dectin-1, suppressing C5a-mediated effector phase of the complement cascade ^69^. These findings support a model in which *TNFRSF13B* diversity in a population establishes a threshold for inflammatory properties of IgG, securing diverse immunologic responses controlling transmission at the cost of excessive inflammation.

IgG glycans, including fucose, sialic acid, and galactose, determine antibody effector functions such as complement activation and antibody-dependent cellular cytotoxicity in complex ways ^53,69–71^. Reduced IgG sialylation and galactosylation enhance complement recruitment ^53,69^ and are clearly and consistently associated with pro-inflammatory diseases in humans^72^. To our knowledge, only a limited number of genetic studies have identified gene polymorphisms associated with fucosylation of plasma proteins ^73,74^. Josipović et al (2019) identified that genetic polymorphisms in *HNF1A* (hepatocyte nuclear factor 1-alpha) can influence plasma protein fucosylation^73^. Lauc et al (2010) identified the rs7159888 SNP of the *FUT8* gene as an effector of plasma protein fucosylation ^74^. Knockdown in the *IKZF1* (Ikaros family zinc finger 1) gene increases IgG fucosylation ^75^.

The question remains, how TNFRSF13b polymorphisms determine glycosylation of IgG. IgG is glycosylated in B cells ^76,77^ and appropriate glycosylation is necessary for structural integrity of IgG ^70^. Glycosylation of IgG by B cells depends on the developmental stage of the cell ^78^. Wang et. al ^79^ showed that plasmablasts early on in immune responses produce less glycosylated IgG, whereas long-lived plasmablasts exhibit elevated IgG glycosylation.

Endothelial cells recycle IgG in a FcRn dependent manner and in this process add sialic acid to IgG ^80^. But since endothelial cells do not express TACI, we hypothesized that the *Tnfrsf13b*-dependent changes in IgG glycosylation are B cell autonomous. Consistent with that idea, we found that CD138+ plasma cells isolated from the spleen of *Tnfrsf13b* mutant mice express less sialyl-transferase ST6GAL1 and less galactosyl-transferase B4GALT1 than plasma cells isolated from WT mice.

Although we have not shown a direct link between TNFRSF13B signaling, transcription, and/or translation of glycosyltransferases, B cells expressing Tnfrsf13b A144E variants have impaired NF-κB signaling, both at baseline and after stimulation with CD40L and IL-4. CD40 synergizes with TNFRSF13B to promote B cell activation and differentiation^40,46^. Hence, by impairing B cell signaling in response to T-cell dependent stimuli, our results suggested that the A144E variant decreases expression of glycosyl-transferases either directly or indirectly due to changing the kinetics of B cell differentiation. Consistent with the latter, we have found that expression of Blimp-1, necessary for plasma cell differentiation and antibody secretion, is decreased in germinal center B cells in Tnfrsf13b A144E variant mice following allogeneic cell immunization^81^.

In summary, the high frequency of *TNFRSF13B* missense alleles in the population is not consistent with a detrimental impact on health. In fact, results in this manuscript argue for a “protective” function of missense alleles that increases effector functions of antibodies, which are key to the clearance of microbes. The *TNFRSF13B-*genotype-dependent enhanced function of antibodies may, however, come at a cost. Consistent with that idea, we found that *TNFRSF13B-*genotype-dependent antibody effector functions are associated with robust inflammatory responses after SARS-CoV-2 and the development of acute respiratory distress syndrome ^82^. The balance between enhanced host defense, possibly decreased transmission, and heightened inflammation might reflect balancing selection that helps to explain the persistence of *TNFRSF13B* mutant alleles in the human population.

## MATERIALS AND METHODS

### Mice

C57BL/6 WT mice were purchased from The Jackson Laboratory (C57BL/6J, IMSR catalog JAX: 000664). Mice harboring bi (A144E/A144E) or monoallelic A144E variants (A144E/WT) are described in this study ^47^. Mice lacking secretory IgA (pIgR-KO) due to deficient polymeric Ig receptor is described here ^83^. All the Tnfrsf13b mutant mice were bred into the C57BL/6 background. Most mice used in the experiment shared same A144E/WT mothers to normalize microbiota in addition to bed-swapping between cages two weeks prior to experiments. Animals of both sexes between 8-20 weeks of age were used.

### Host-adapted *C. rodentium* infection

The kanamycin-resistant (Km-resistant) WT *C. rodentium* strain DBS120 (pCRP1:Tn5) was gifted by David Schauer, Massachusetts Institute of Technology, Cambridge, Massachusetts, USA ^84^. The *C. rodentium* expressing GFP was a gift from Bruce Vallance, University of British Columbia, Vancouver, Canada ^85^. For the host-adapted *C. rodentium* infection, C57BL/6 WT mice are infected with 10^8^ CFU of overnight-LB culture *C. rodentium* DBS120 by oral gavage. 7 days post-infection, feces from infected C57BL/6 WT mice are collected and homogenized with 5 mL of PBS. The homogenized mixture is passed through a 70 μM cell strainer, and the resulting fecal solution containing the host-adapted *C. rodentium* is centrifuged at 3000 rpm for 15 minutes and reconstituted with 5 mL of fresh PBS. 200 μL of the slurry containing approximately 10^7^ CFU of host-adapted *C. rodentium* is used to oral gavage a new cohort of WT and Tnfrsf13b variant A144E mice. To have a better estimate of the amount of feces from 7-day lab-strain *C. rodentium*-infected WT mice needed for the host-adapted *C. rodentium* infection with 10^7^ CFU, CFU/mg of feces is determined at 6 days post-infection.

### CFU quantification

At the designated time points during infection, ∼50-75 mg of feces was collected, weighed, and suspended in 500 μL of PBS. The fecal pellets are then homogenized at 2500 rpm for 5 minutes using a Disruptor Genie (Scientific Industry). Optimally diluted homogenized fecal mixture is plated on kanamycin-containing MacConkey agar plates and incubated overnight at 37 °C. The number of colonies were determined, and the number of bacteria per 1 mg was calculated. Infection was considered cleared when less than 10^1^ CFU was detected in the feces suspension.

### *C. rodentium* Intimin purification

The intimin expression strain was gifted by Dr. Gabriel Nunez. Briefly, the construction plasmid contains the extracellular C-terminal 385 amino acids of intimin-beta which is cloned into the expression vector pQE32. The glycerol stock of the intimin expression strain was cultured overnight in 10 mL of LB broth supplemented with 50 μg/mL of ampicillin and kanamycin at 30 °C with 225 RPM agitation. This culture was used to scale to 1 L of LB broth with the same conditions as above. The next day, 500 μL of 20 μM IPTG was added to this culture which was then scaled to 2 L total and incubated again overnight in the same conditions. The next day, the culture was centrifuged at 5000 rpm for 30 minutes to form a pellet. To purify the intimin protein from this bacterial pellet, a Ni-NTA Fast Start Kit (Qiagen 30600) was used according to directions to lyse the bacteria and purify the protein. The purified intimin protein was then concentrated with Amicon Ultra-15 Filters Units (Sigma Aldrich UFC9010D). This intimin was then quantified by Bradford assay.

### *C. rodentium* LPS purification

The DSB120 *C. rodentium* strain was grown as previously described. The *C. rodentium* overnight culture was centrifuged at 5000rpm for 30 minutes to form a pellet. This bacterial pellet was then lysed and purified using an LPS extraction kit (Abcam ab239718). The LPS in the purified product was then quantified using a Total Carbohydrate Assay Kit (Abcam ab155891).

### Quantification of Total Ig

MaxiSorp ELISA plates (Thermo Fisher Scientific 442404) were coated overnight with either goat anti– mouse Ig diluted 1:250 (4ug/mL). The plates were then blocked for a minimum of 1 hour. Serial dilutions of samples from immunized mouse serum or fecal supernatant were added to the coated wells. Then, bound Ig was detected using a goat anti-mouse IgG HRP, goat anti-mouse IgA HRP, or goat anti-mouse IgM HRP (SouthernBiotech, Catalog #1036-05, 1040-05, or 1021-05 respectively). To obtain OD405 nm readings, 1-Step™ ABTS substrate solution (Thermo Fisher Scientific 37615) was used to react with bound HRP. These absorbance values were read using a plate reader machine. All plate readings were recorded at 405 nm. Washings of plates three times with PBS were conducted by automated plate washer machine in between incubation steps.

### Antigen-specific Ig Quantification

MaxiSorp plates were coated with 1ug of intimin or LPS per well overnight at 4C. For heat inactivated *C. rodentium* specific antibodies, overnight cultured *C. rodentium* is heat-inactivated at 56C for 1 hour and 50uL of it was used to coat each well overnight at 4C. Serial dilution of 1:100 mice serum or fecal supernatant was added to the coated wells. Remaining ELISA steps are completed following the protocol described above.

### *C. rodentium* specific IgG Avidity

Amount of *C. rodentium* specific IgG is determined in a separate ELISA following the protocol described above. Serially diluted *C. rodentium* specific IgG, starting from 0.75ug, is incubated with log-phase *C. rodentium* expressing GFP for 30 minutes at 4C. Following incubation, the bacteria is washed three times with PBS. 1uL of goat-anti-mouse IgG F(ab’) PE (Thermofisher, Catalog# A10543) is added to the bacteria-IgG complex and incubated at room temperature for 30 minutes. Following three washes with PBS, MFI of IgG bound to *C. rodentium* was determined using BD Fortessa Flow Cytometry Analyzer.

### Quantification of IgG sialyation, galactosylation, and core fucosylation

IgG was purified from mouse serum using the Melon Gel IgG Spin Purification Kit (Thermo Fisher, Cat# 45212). Following purification, IgG purity was confirmed by performing ELISA and SDS-PAGE to check for the presence of IgM and IgA. The purified IgG was quantified by ELISA, and 25 µg of IgG was used to coat each well in a serial dilution manner of a MaxiSorp 96-well plate and the plates were incubated overnight at 4°C. The following day, the plates were washed three times with 1x PBS and then blocked by adding 200 µL of 1x Carbo-Free Blocking Solution (Vector Laboratories. SKU# SP-5040-125) per well for 1 hour at room temperature. After blocking, the samples were incubated with a divalent ion solution (1 mM CaCl₂, 1 mM MgCl₂, 1 mM MnCl₂) for 15 minutes at room temperature to optimize lectin-IgG binding. Biotinylated lectins RCA-1, SNA-EBL, LCA (Vector Laboratories, SKU# B-1085-1, B-1305-2, B-1045-5) were diluted to 5 µg/mL in 1x Carbo-Free Blocking Solution, and 100 µL of the diluted lectins was added to each well, followed by incubation for 1 hour at room temperature. Signal detection was carried out by adding 6.25ng of Streptavidin-HRP (Thermo Fisher, Cat# N100) to each well and incubated at room temperature for 45 minutes. For color development, 50µL of 1-step ABTS solution (Thermo Fisher, Cat# 37615) was added to each well, and absorbance was measured at 405nm with a Biotek luminescence plate reader. To normalize ELISA absorbance values across multiple plates for inter-plate variability, absorbance values from all ELISA plates are combined into a single matrix and global mean and standard deviation of the full dataset is calculated. For each absorbance value x, z-score was calculated using the formula, Z= (x – global mean)/standard deviation.

### Glycosyltransferases expression analysis

Naïve mice were sacrificed in accordance with approved animal protocols at the University of Michigan or Case Western Reserve University. Spleens were harvested, and single-cell suspensions were generated by passing the tissue through a 70-µm cell strainer into sterile 50-mL conical tubes. Red blood cells were lysed using ACK lysis buffer for 4 minutes at room temperature, followed by washing with RPMI medium. Cells were counted, and 4 million cells were plated in 4 mL of B-cell culture medium. The B-cell medium consisted of RPMI supplemented with 10% heat-inactivated FBS, L-glutamine, penicillin-streptomycin, and 500 µL β-mercaptoethanol (BME) per 500 mL of medium. After 72 hours of culture, cells were collected and centrifuged at 1500 rpm for 10 minutes. Supernatants were harvested and stored for subsequent lectin analysis as described above. Cell pellets were resuspended in cell suspension buffer for flow cytometry. For surface staining, cells were incubated with 1 µL anti-mouse CD19 APC (BioLegend, Cat#152410) and 1 µL anti-mouse CD138 PerCP/Cy5.5 (BioLegend, Cat#142510) for 30 minutes at room temperature. After three washes, cells were fixed and permeabilized using the eBioscience Intracellular Fixation and Permeabilization Buffer Set (Invitrogen, Cat#88-8824-00) according to the manufacturer’s instructions. For intracellular staining, cells were incubated for 30 minutes at room temperature with 1 µL of either anti-mouse ST6Gal1 (Proteintech, Cat#14355-1-AP) or anti-mouse B4GALT1 (Proteintech, Cat#55448-1-AP). Following washing, goat anti-rabbit IgG PE (Southern Biotech, Cat#4050-09) was used as a secondary antibody and incubated for 30 minutes at room temperature. Cells were analyzed using a BD LSR Fortessa flow cytometer.

### NF-κb signaling in CD19+ B cells

Spleens were harvested from mice, and single-cell suspensions were prepared as described above. A total of 1 × 10 cells were plated in 2 mL of B-cell culture medium and rested for 45 minutes at 37°C. Cells were stimulated with 1.5 µg CD40L and 0.30 µg IL-4 per well for 15 minutes at 37°C. Following stimulation, cells were collected and washed twice with cell suspension buffer. Cells were fixed and permeabilized using the BD Pharmingen Transcription Factor Buffer Set (BD, Cat#562575) according to the manufacturer’s instructions. Cells were then stained with 1 µL anti-mouse CD19 APC (BioLegend, Cat#152410) and 1 µL anti-mouse phospho-NF-κB p65 (Ser536) (Invitrogen, Cat#MA5-15160) for 30 minutes at room temperature. After washing, goat anti-rabbit IgG PE (Southern Biotech, Cat#4050-09) was used as a secondary antibody and incubated for 30 minutes at room temperature. Cells were analyzed using a BD LSR Fortessa flow cytometer.

### In vivo complement deposition analysis

Mice were infected with GFP-expressing *C. rodentium* according to the infection protocol described above. Ten days post-infection, mice were sacrificed, and fecal pellets were collected from the proximal colon. Fecal pellets were homogenized in PBS, and the homogenate was sequentially passed through a 70-µm cell strainer followed by a 40-µm cell strainer to remove debris. The filtrate was centrifuged, and the bacterial pellet was resuspended in fresh PBS for flow cytometry analysis. Isolated bacteria were stained with one of the following antibodies: anti-mouse C1q (biotinylated; Invitrogen, Cat#MA1-40312), anti-mouse C3d (Bioss, Cat#BS-4877R), anti-mouse C3/C3b/iC3b PE, or anti-mouse IgG F(ab’)₂ PE (Thermo Fisher Scientific, Cat#A10543). When required, secondary detection was performed using either goat anti-rabbit IgG PE (Southern Biotech, Cat#4050-09) or streptavidin-PE (Invitrogen, Cat#12-4317-87). For flow cytometric analysis on a BD LSR Fortessa, cultured GFP-expressing *C. rodentium* was first used to establish the gating strategy and define the GFP-positive bacterial population. This gating strategy was then applied to identify GFP-positive bacteria isolated from infected mouse feces.

### Measuring free serum C3

Serum C3 was quantified from infected mouse blood using a Mouse Complement C3 ELISA kit (abcam ab157711) following the manufacturer’s instructions. Briefly, serum was diluted at 1:250,000 for this analysis. Diluted serum is incubated with capture and detection antibody cocktail reagent in the 8 coated well strips provided in the kit. Following 1.5-hour incubation at room temperature, wells were washed with provided wash buffer three times then developing solution was used to obtain absorbance values at 405nm using a Biotek plate reader.

### Quantification of intimin specific IgG-complement C1q interaction

MaxiSorp plates were coated with 1 µg/well of purified intimin overnight at 4°C. Following coating, plates were blocked with 2% BSA solution for one hour. Then serial dilution starting from 1 ug of intimin-specific IgG (determined separately in intimin-IgG ELISA) was added per well and incubated at room temperature for one hour. Following incubation with intimin protein and purified IgG, 1 µg of recombinant purified human C1q protein (Complement Technology, Catalog # A099) in PBS with 1mM of CaCl2, and 1mM of MgCl2 was added to each well and incubated for 1 hour at room temperature. After 1 hour incubation with human C1q, plates were washed 3 times with 1x PBS and anti-human C1q biotinylated (ThermoFisher, MA1-40312) was added to each well in the dilution of 1:50. Purified human C1q was used as a complement source as it is more robust than mouse C1q but still has high genetic similarity ^86^. After one hour incubation at room temperature, 6.25 ng of HRP-conjugated Streptavidin-HRP (ThermoFisher, N100) was added per well for color development using 50µL of 1 step ABTS solution (Thermo Fisher, Cat# 37615). Absorbance values were measured at 405 nm using Biotek Luminescence Plate Readers.

### Measuring *C. rodentium* specific IgG sialylation

Overnight cultured *C. rodentium* expressing GFP was blocked with 1x Carbo-Free Blocking Buffer (Vector Laboratories. SKU# SP-5040-125) for 30 minutes at 4C. 0.2 ug of *C. rodentium* specific IgG (determined in a separate ELISA) was incubated with the bacteria for 30 minutes at 4C. Following 3 PBS washes, 100uL of diluted SNA-EBL (5 µg/mL) was added to the samples and incubated for 45 minutes at room temperature. After incubation, 1uL of streptavidin-PE for each sample (Invitrogen, Cat#12-4317-87) was used for detection on BD LSD Fortessa.

### Determining C5b-9 deposition on GFP expressing *C. rodentium*

Log-phase *C. rodentium* expressing GFP was blocked with 2% BSA solution for 30 minutes at 4C. 0.25 ug of *C. rodentium* specific IgG (determined in a separate ELISA) was incubated with the bacteria for 30 minutes at 4C. Following 3x PBS washes, 50uL of human antibody-depleted serum (Pel-Freez, Cat: 34041-1) was added to the samples and incubated for 30 minutes at 37C with shaking. Following incubation at 37C, 1µL of anti-C5b-9 (ae11) antibody (concentrated to 0.5mg/mL) (Santa Cruz, Cat: sc-58935) was used to detect C5b-9 deposition on GFP expressing *C. rodentium*. For flow cytometry use, anti-C5b-9 antibody is conjugated with PE / R-Phycoerythrin using abcam PE / R-Phycoerythrin Conjugation Kit -Lightning-Link (Cat: ab102918). Samples without IgG or serum complement source were also included to account for background.

## ACKNOWLEDGMENTS

We would like to acknowledge Dr. Gabriel Nunez (University of Michigan) for providing key reagents for the research and critically read the manuscript. We are thankful for David Shauer for the gift of *C. rodentium* strain DBS120. Also, for Bruce Vallance and Nobuhiko Kamada.

**Grants to acknowledge are the following:** AI 173950, AI 159219, GM 140359 to MC and JLP

We acknowledge the **Richard J. Fasenmyer Professorship in Immunopathogenesis to MC, Case Western Reserve University and the School of Medicine**

**This research was supported by the Cytometry & Imaging Microscopy Shared Resource of the Case Comprehensive Cancer Center (P30CA043703)**

## AUTHOR CONTRIBUTIONS

Conceptualization, LN, JLP, MC; methodology, LN, IC, NG, SS, IKL, AH, CCP, CK, MB, TJP, JLP, MC; Investigation, LN, IC, NG, SS, IKL, AH, CCP, CK, MB, TJP, JLP, MC; writing—original draft, LN, IC, MC; writing—review & editing, LN, IC, CK, VPR, MC; funding acquisition, LN, JLP, MC; resources, INITIALS; supervision, LN, JLP, MC.

## SUPPLEMENTAL INFORMATION

**Document S1. Figures S1–S5,**

