## Supplementary material for "Tnfrsf13b Variants Enhance IgG-Mediated Complement Responses to Promote Bacterial Clearance": Suplemental material

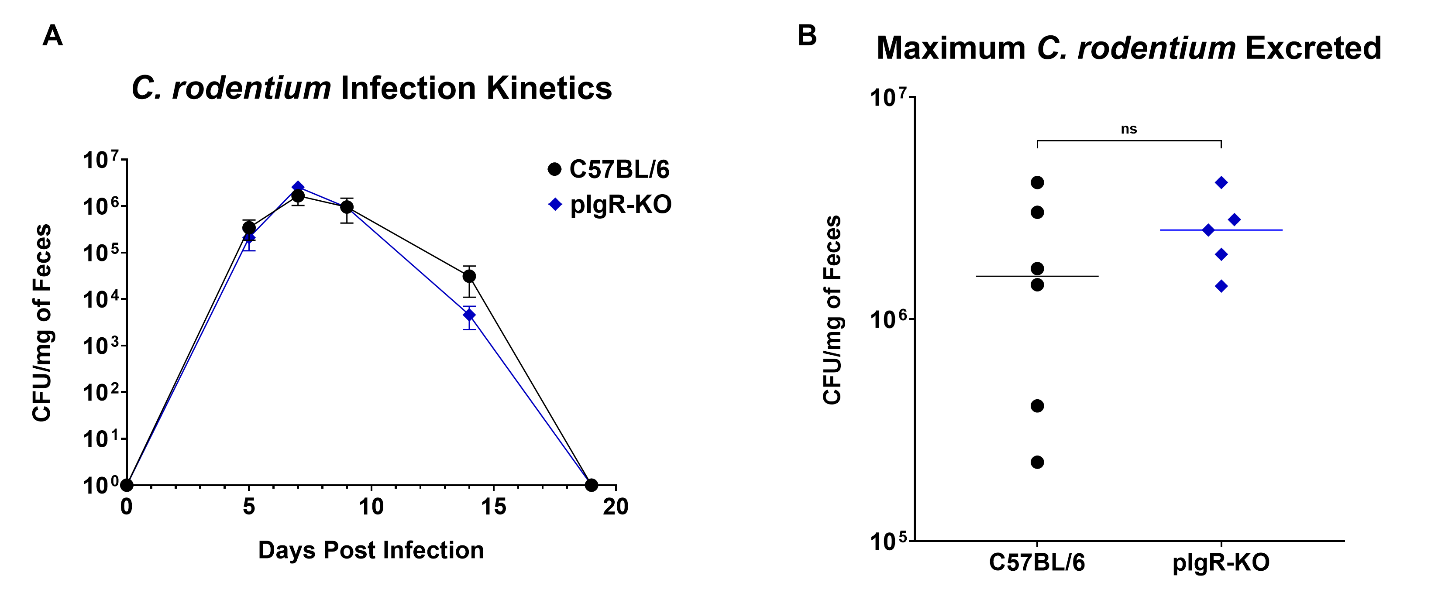

**Figure S1.** **Host-adapted *C. rodentium* infection in pIgR-KO mice, related to Figure 1**

pIgR-KO mice do not have polymeric immunoglobulin receptor for translocation of IgA to epithelial surfaces. (A) Graph depicts the kinetics of *C. rodentium* infection quantified by the mean number of viable *C. rodentium* in feces (expressed in log_10_ CFU/mg of feces) at various times after infection. (B) Peak bacterial burden (maximum log₁₀ CFU/mg feces) observed during infection for each genotype. WT (n = 6), pIgR-KO (n = 5). Statistical analysis was performed using ordinary Mann-Whitney Test in GraphPad Prism v10.4.1.

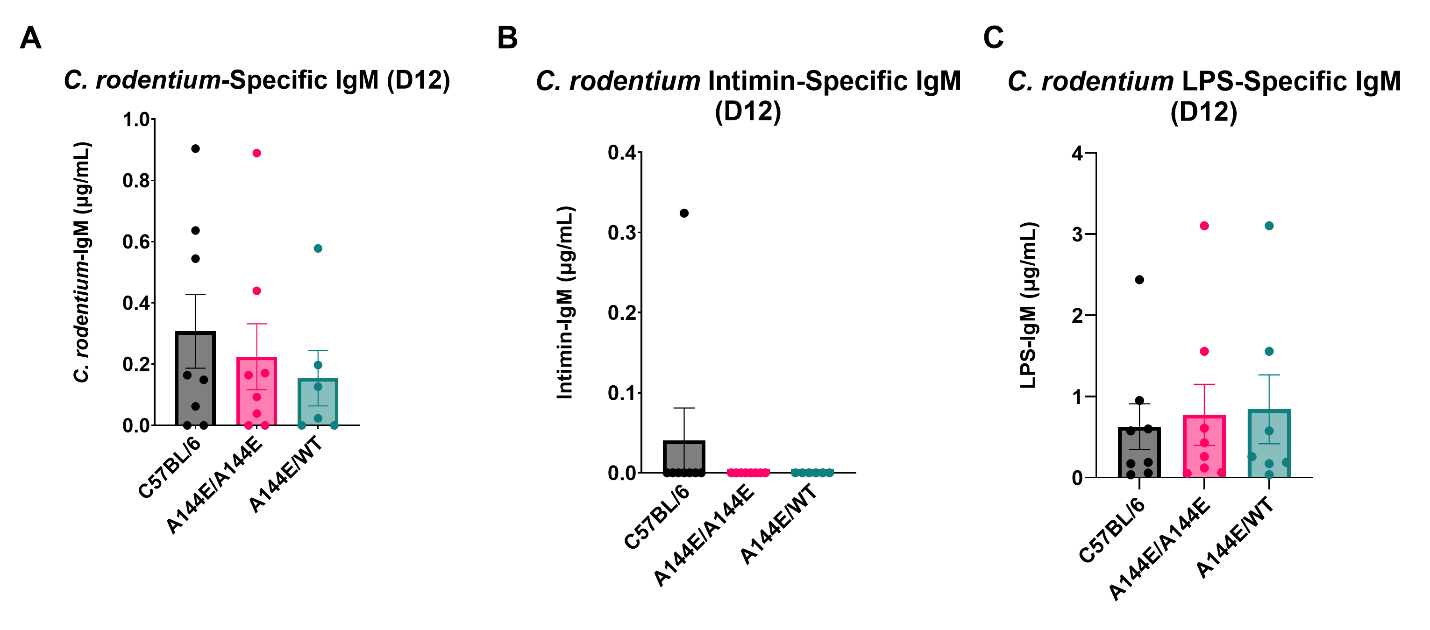

**Figure S2: *C. rodentium* specific IgM antibodies in *Tnfrsf13b* mutant mice, related to Figure 2**

Serum from infected mice were collected at indicated time points during infections. Amounts of (A) intimin-specific IgM, (B) LPS-specific IgM and (C) heat-inactivated *C. rodentium* specific IgM, were measured. WT (n = 8), A144E/A144E (n = 8), A144E/WT (n = 6). Bar graphs show mean IgG levels. Statistical comparisons between genotypes are made with two-way ANOVA followed by Fisher’s LSD test in GraphPad Prism v10.4.1.

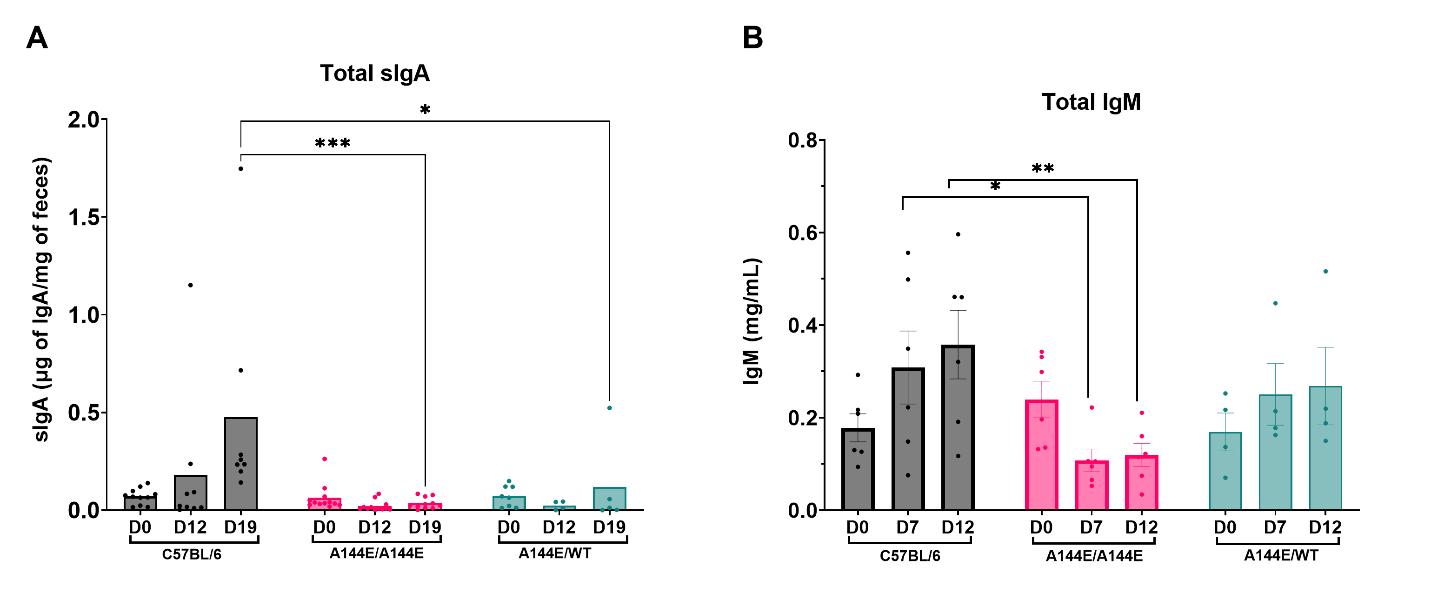

**Figure S3: Total secretory IgA (sIgA) and total IgM antibodies in *Tnfrsf13b* mutant mice, related to Figure 2**

Fecal supernatant or serum from infected mice were collected at indicated time points during infections. Amounts of (A) total sIgA; WT (n = 12), A144E/A144E (n = 12), A144E/WT (n = 8) and (B) total IgM; WT (n = 6), A144E/A144E (n = 6), A144E/WT (n = 4) were measured. Bar graphs show mean Ig levels. Statistical comparisons between genotypes are made with two-way ANOVA followed by Fisher’s LSD test in GraphPad Prism v10.4.1.

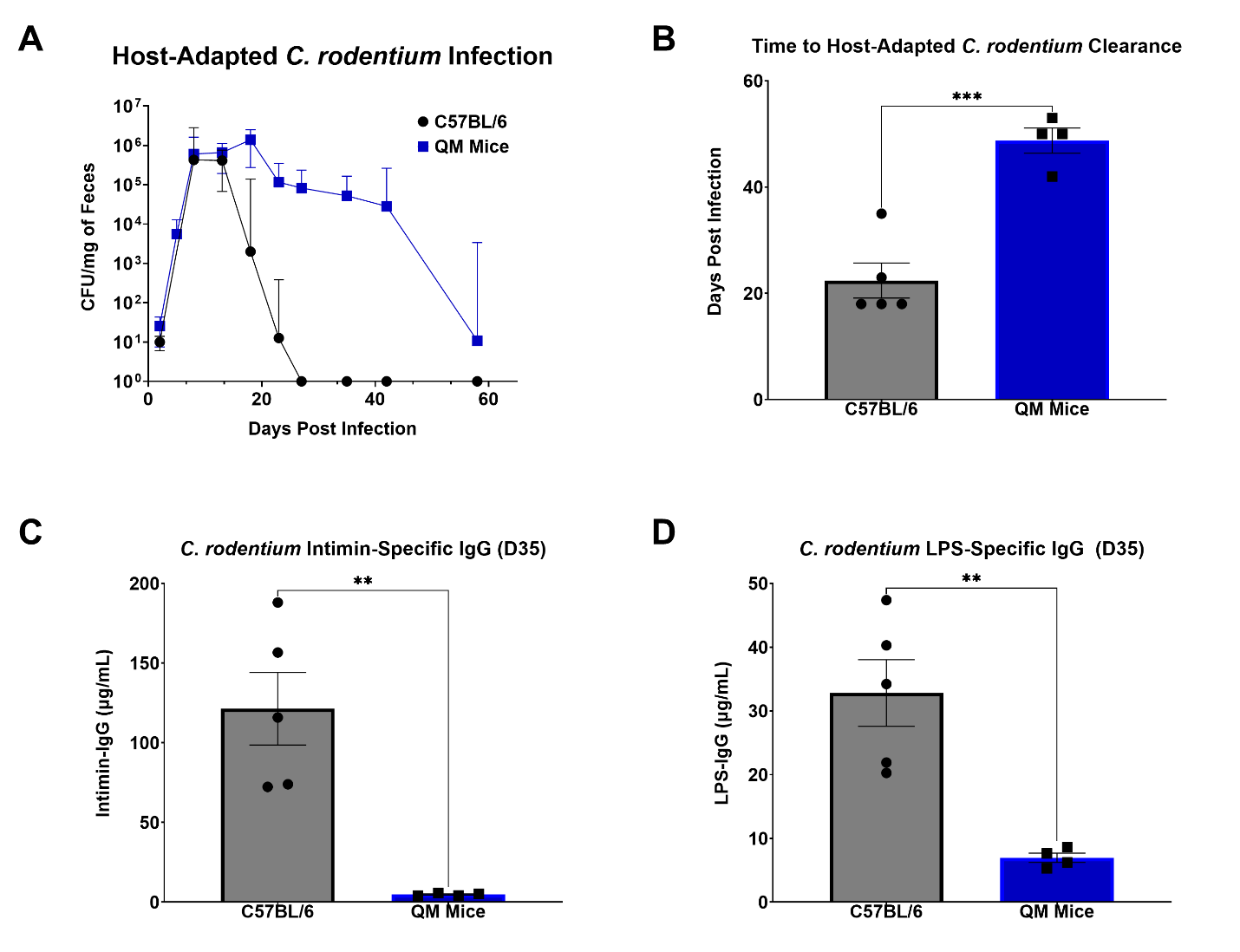

**Figure S4.** **Host-adapted *C. rodentium* infection in QM mice, related to Figures 1 & 2**

Quasi-monoclonal mice (QM) are unable to make antigen-specific antibodies as their B cell repertoire is limited to hapten-4-hydroxy-3-nitrophenylacetate. (A) Graph depicts the kinetics of *C. rodentium* infection quantified by the mean number of viable *C. rodentium* in feces (expressed in log_10_ CFU/mg of feces) at various times after infection. (B) Bar graph shows the average time to clear bacteria, time to clearance is defined by when detected CFU is below 10^1^ CFU/mg of feces. On day 35 post infection, serum was collected and (C) intimin-specific IgG and (D) LPS-specific IgG were measured. WT (n= 5); QM mice (n= 4). Statistical analysis was performed using ordinary Mann-Whitney Test in GraphPad Prism v10.4.1.

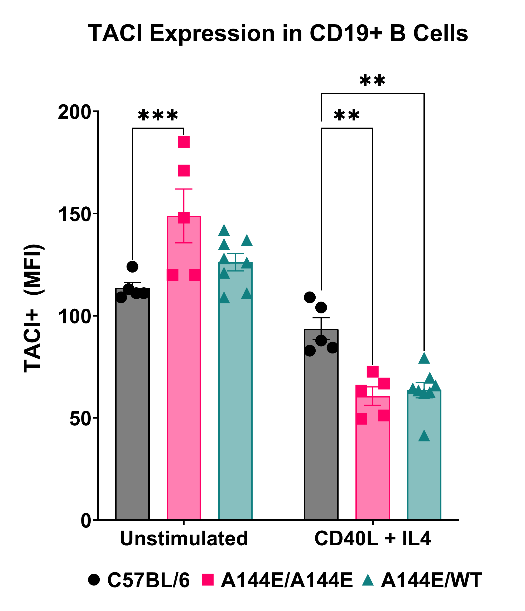

**Figure S5. TACI expression in CD19+ B cells, related to Figure 4**

Splenocytes were isolated from WT mice (n=5), A144E/A144E mice(n=5), and A144E/WT mice(n=8), and stimulating them with CD40L+ IL4. Unstimulated and stimulated cells after 72-hour culture in B cell media were fixed, permeabilized, and stained with anti-CD19 antibody and antibody for TACI. Representative MFI distribution curves and corresponding quantification (bar graphs) for WT (n=5), A144E/A144E (n=5), and A144E/WT (n=8) are shown. Statistical comparisons between genotypes are made with two-way ANOVA followed by Fisher’s LSD test in GraphPad Prism v10.4.1.
